# Label-free and non-destructive pathology of human lung adenocarcinomas with ultraviolet single-plane illumination microscopy

**DOI:** 10.1101/2023.01.02.522477

**Authors:** Yan Zhang, Bingxin Huang, Lei Kang, Victor T. C. Tsang, Jiajie Wu, Claudia T. K. Lo, Terence T. W. Wong

## Abstract

Lung cancer is one of the leading causes of cancer death worldwide. The diagnosis of lung cancer based on the analysis of formalin-fixed and paraffin-embedded (FFPE) tissues is laborious and time-consuming, failing to guide surgeons intraoperatively. Here we proposed a rapid histological imaging method, termed microscopy with ultraviolet single-plane illumination (MUSI), to enable rapid *ex*-or *in*-*vivo* imaging of fresh and unprocessed tissues in a label-free and non-destructive manner. The MUSI system allows surgical specimens with large irregular surfaces to be screened at a speed of 0.5 mm^2^/s with a subcellular resolution, which is sufficient to provide immediate feedback to surgeons and pathologists for intraoperative decision-making. We demonstrate that MUSI can differentiate between different subtypes of human lung adenocarcinomas, revealing diagnostically important features that are comparable to the gold standard FFPE histology. As an assistive imaging platform, MUSI could facilitate the development of precise image-guided surgery and revolutionize the current practice in surgical pathology.

## 1. Introduction

Lung cancer is one of the leading causes of cancer mortality worldwide, with an estimated 2.2 million new cancer cases and 1.8 million deaths in 2020 [1]. 85% of the diagnosed cases fall into the category of non-small cell lung cancer (NSCLC), with a 5-year survival rate of 21% [2]. Adenocarcinoma is the most common type of NSCLC, which is usually developed with a mixture of histologic subtypes. Since 40% of NSCLCs will have metastasized beyond the lungs by the time it is diagnosed, early detection of NSCLC is crucial to improving the treatment performance. Currently, surgery to remove the affected tissue or tumor is the most curative treatment option for early-stage NSCLCs. Histopathology has remained the gold standard for surgical margin assessment (SMA) for decades. However, routine pathological examination based on formalin-fixed and paraffin-embedded (FFPE) tissues is laborious and time-consuming, failing to provide immediate on-site feedback to surgeons and pathologists for intraoperative decision-making. Although frozen section can serve as a rapid alternative to FFPE, it still requires a turnaround time of 20 to 30 minutes during surgery [3]. Besides, the frozen section is subjected to inadequate sampling of resection margins, and freezing artifacts are inevitable when dealing with lipid-rich tissues. All these factors will affect the histopathological interpretation and diagnostic accuracy [4].

Recent advances in optical microscopy enable rapid and slide-free imaging of thick fresh tissues, holding great promise to streamline the current practice in FFPE histology [5]. Imaging modalities based on exogenous fluorophores, including fluorescence confocal microscopy [6]–[8], light-sheet microscopy [9], [10], structured illumination microscopy [11], [12], and microscopy with ultraviolet (UV) surface excitation [13], [14], can provide sufficient sampling of large resection margins within a point-of-care timeframe, providing highly specific cellular features for pathological diagnosis. However, they pose a threat to intraoperative procedures as some new fluorescent contrast agents might introduce toxicities to patients. Besides, the staining process may interfere with the subsequent molecular assays such as fluorescence *in situ* hybridization and DNA/RNA sequencing [10]. In contrast, imaging techniques based on intrinsic contrast mechanisms are highly desired in modern clinical settings. For instance, with the nature of deep penetration and non-invasiveness, optical coherent tomography [15], [16] and reflectance confocal microscopy [17], [18] have been successfully translated in the area of ophthalmology and dermatology for years. However, the cellular contents provided by these reflectance-based methods are relatively limited within internal organs. Besides, photoacoustic microscopy [19], [20] can spectrally probe different molecular targets with intrinsic absorption contrast, showing promising results in vascular imaging and breast cancer screening. In addition, nonlinear processes [21]–[24], including coherent Raman scattering, multiphoton absorption, and harmonic generation, can achieve high-resolution and label-free visualization of a variety of biological processes in an unperturbed and non-destructive way. However, these methods still face challenges in screening large-scale specimens within a short diagnostic timeframe due to the requirement of sequential beam scanning.

Our recently proposed method, termed computational high-throughput autofluorescence microscopy by pattern illumination (CHAMP) [25], enables rapid, high-resolution, and label-free histological imaging of thick and unprocessed tissues, particularly favoring the applications of intraoperative SMA where immediate feedback should be provided to surgeons for optimal adjuvant treatment. However, the depth-of-focus (DOF) of CHAMP is restricted to 80 μm, which is not sufficient to accommodate large surface irregularities presented in manually-cut tissues, causing resection margins to come in and out of focus during imaging. Besides, CHAMP achieves optical sectioning by leveraging the shallow penetration of deep-UV light into scattering biological tissues. This varies between different types of tissues [14], such that the CHAMP images could exhibit slight deviations from slide-based FFPE histology. To overcome these limitations, here we propose a new imaging method, termed microscopy with ultraviolet single-plane illumination (MUSI). Similar to light-sheet microscopy, MUSI rejects out-of-focus fluorescence by illuminating the specimen with a thin sheet of light and detects the resulting signal from an orthogonal direction. The dual-axis configuration of MUSI decouples the illumination from detection beam paths, overcoming the inherent tradeoff between long DOF and high spatial resolution in a conventional epi-illumination setup. In addition, rich endogenous fluorophores [26], including reduced nicotinamide adenine dinucleotide (NADH), structural proteins (e.g., collagen and elastin), aromatic amino acids (e.g., tryptophan, tyrosine), and heterocyclic compounds (e.g., flavins, lipopigments), naturally form a fundamental contrast mechanism with deep-UV excitation. By taking advantage of this intrinsic fluorescence, MUSI enables label-free and non-destructive imaging of fresh and unprocessed tissues at an imaging speed of 0.5 mm^2^/s, achieving a spatial resolution of 2 μm (lateral) and 2.8 μm (axial) with a long DOF up to ∼200 μm. This guarantees high-quality histology-like images can be obtained at the surface of irregular surgical tissues. We demonstrate that MUSI can differentiate between different subtypes of human lung adenocarcinomas (*n* = 15), revealing diagnostically important features that are comparable to the gold standard histology. Our results suggested that MUSI has great potential as an assistive diagnostic tool that can be used by surgeons and pathologists for intraopertive decision-making.

## 2. Methods

### 2.1. Preparation of biological tissues

For *ex-vivo* imaging, the internal mouse organs, including the liver, brain, kidney, lung, spleen, skin, muscle, and tongue, were harvested immediately after the mice (C57BL/6 type) were sacrificed. After extraction, these organs were kept intact or manually cut into 3-to 5-mm-thick tissue slabs for imaging without any further processing. For *in-vivo* imaging, the mice were supinated on the sample holder and heavily anesthetized with 3% isoflurane during experiments. To expose the targeted brain and kidney tissues for imaging, a 5-mm by 5-mm cranial window was made on the skull and a small incision was made on the left abdomen. For human lung tissues, the specimens were obtained from lung cancer patients who underwent curative lung cancer surgery at the Queen Mary Hospital. Following lung lobectomy, the cancer tissues were cut with a scalpel from the resected lobe, and subsequently formalin-fixed and transported to the lab for imaging. After imaging, all the specimens were histologically processed with a standard protocol to obtain the hematoxylin and eosin (H&E)-stained images. Specifically, the specimens were fixed in 4% neutral-buffered formalin at room temperature for 24 hours, and processed for dehydration and infiltration by a tissue processor (Revos, Thermo Fisher Scientific Inc.) for 12 hours. After that, the specimens were paraffin-embedded and subsequently sectioned into 4-μm-thick tissue slices by a microtome (RM2235, Leica Microsystems Inc.). Finally, the sectioned thin tissue slices were mounted on glass slides, stained by H&E, and imaged by a digital slide scanner (NanoZoomer-SQ, Hamamatsu Photonics K.K.) to generate the corresponding histological images. All animal experiments were carried out in conformity with the guidelines and protocols approved by the Health, Safety and Environment Office (HSEO) of the Hong Kong University of Science and Technology (HKUST) (license number: AH18038). All human experiments were carried out in conformity with a clinical research ethics review approved by the Institutional Review Board of the University of Hong Kong/Hospital Authority Hong Kong West Cluster (HKU/HA HKW) (reference number: UW 20-335). Informed consent was obtained from all lung cancer tissue donors.

### 2.2. 4T1 cell allograft mouse model

#### 2.2.1. 4T1 cell culture

The mouse breast cancer 4T1 cell line was purchased from American Type Culture Collection (ATCC, Manassas, VA, USA). The cells were grown in RPMI 1640 medium supplemented with 10% fetal bovine serum and 1% penicillin-streptomycin solution and cultured at 37°C in a 5% CO_2_ incubator. Subconfluent 4T1 cells were trypsinated and resuspended in phosphate-buffered saline (PBS). 10 μL of the resuspension was mixed with trypan blue, and the number of 4T1 cells was counted using an automated cell counter (Countess™ 3 FL Automated Cell Counter, Thermo Fisher Scientific). A cell resuspension with a final cell density of 1 × 10^6^ cells/ml was prepared in PBS.

#### 2.2.2. 4T1 orthotopic allograft

6–8 weeks-old female Balb/c mice were used to generate the 4T1 cell allograft mouse model. Hair around the injection site, the left abdominal mammary gland, was removed using hair clippers and was sterilized with three alternating swabs of betadine and 70% ethanol. 1 ml tuberculin syringe with a 27 gauge needle was used to inject 200 μL of 4T1 cells (2 × 10^5^ cells) subcutaneously. A palpable primary tumor was developed and observed after one week. The mice injected with the cells were euthanized by carbon dioxide asphyxiation three weeks after injection of 4T1 cells. Organs and tissues were then harvested.

### 2.3. Configuration of the MUSI system

The setup of the MUSI system shares similarities with the previously reported open-top light-sheet configurations [9], [27], [28], which offer great flexibilities to manipulate large and thick samples without the use of immersion objectives. As shown in Fig. 1a, the system consists of two orthogonal beam paths under the specimen that sits at 45° with respect to the optical axes. A customized 20-mm-diameter UV-transparent quartz hemisphere (*n* = 1.45, Worldhawk Optics Ltd.) is held beneath a quartz-bottom sample holder (1-mm thickness) to mitigate imaging aberrations induced by oblique detection at the air-holder interface [9]. Both illumination and detection paths focus perpendicularly through the hemisphere and conjugate at the sample holder. A thin layer of UV-transparent oil (#56821, Sigma-Aldrich) is applied in the space between the holder and the bottom surface of the tissue to achieve optimal index matching. For the illumination arm, a nanosecond UV pulsed laser is used as the excitation source (WEDGE HF 266 nm, Bright Solutions Srl.), which is spectrally filtered by a bandpass filter, F1 (FF01-300/SP-25, Semrock Inc.), and expanded by a pair of lenses, L1 and L2 (LA4647-UV and LA4874-UV, Thorlabs Inc.). Then the beam is propagated through an adjustable slit aperture, SA (VA100C, Thorlabs Inc.), outputting a beam with a size of 2 mm × 6 mm, which is subsequently focused by a UV cylindrical lens, CL1 (LJ4395RM, *f* = 100 mm, Thorlabs Inc.), and generates a 2-mm-wide Gaussian light sheet with a waist radius (*w*_0_) of 2.8 μm and a DOF (2*Z*_R_) of 190 μm (Fig. S1). For the detection arm, the excited intrinsic fluorescence is collected by an achromatic UV objective lens, OL (LMU-5X-NUV, NA = 0.12, Thorlabs Inc.), filtered by a long pass filter, F2 (BLP01-325R-25, Semrock Inc.), and subsequently refocused by an infinity-corrected tube lens, TL (TTL180-A, Thorlabs Inc.), transmitted through an additional low-power cylindrical lens, CL2 (*f* = 2000 mm, Worldhawk Optics Ltd.), and finally imaged by an sCMOS camera (PCO edge 4.2, 2048 × 2048 pixels, PCO Inc.). The specimen is raster scanned by a 3D high-speed motorized stage (L-509.20SD00, PI miCos GmbH) with a maximum traveling range of 5 cm. The specimen is translated through the static light sheet at a constant velocity of 0.25 mm/s along the primary scanning direction (x-axis in Fig. 1b), and the images are recorded at 250 frames/s with a sampling pitch of 1 μm/pixel at the camera plane. The image height can be adjusted according to the surface irregularities of the imaged specimen, with a maximum tolerance of ∼200 μm in the current design. Followed by the primary scanning, the specimen was translated laterally (y-axis in Fig. 1b) at an interval of 1.8 mm, causing a 10% overlap between adjacent image stripes for stitching large field-of-views (FOVs). The raw images were stored at 16-bit depth and transferred through a CameraLink interface at a streaming rate of 2000 pixels (w) × 200 pixels (h) × 2 Byte × 250 frames/s = 0.2 GB/s to a local workstation equipped with high-speed solid-state disks (970 EVO Plus, SAMSUNG Inc.). The image acquisition and stage scanning were synchronized by our lab-designed LabVIEW software (National Instruments Corp.). The design of MUSI allows label-free and slide-free imaging of fresh and unprocessed tissues at an imaging speed of 0.5 mm^2^/s with an in-plane resolution of ∼2 μm (Fig. S2), producing histology-like images with sufficient anatomic features that can be easily interpreted by pathologists for intraoperative decision-making.

**Figure 1.**
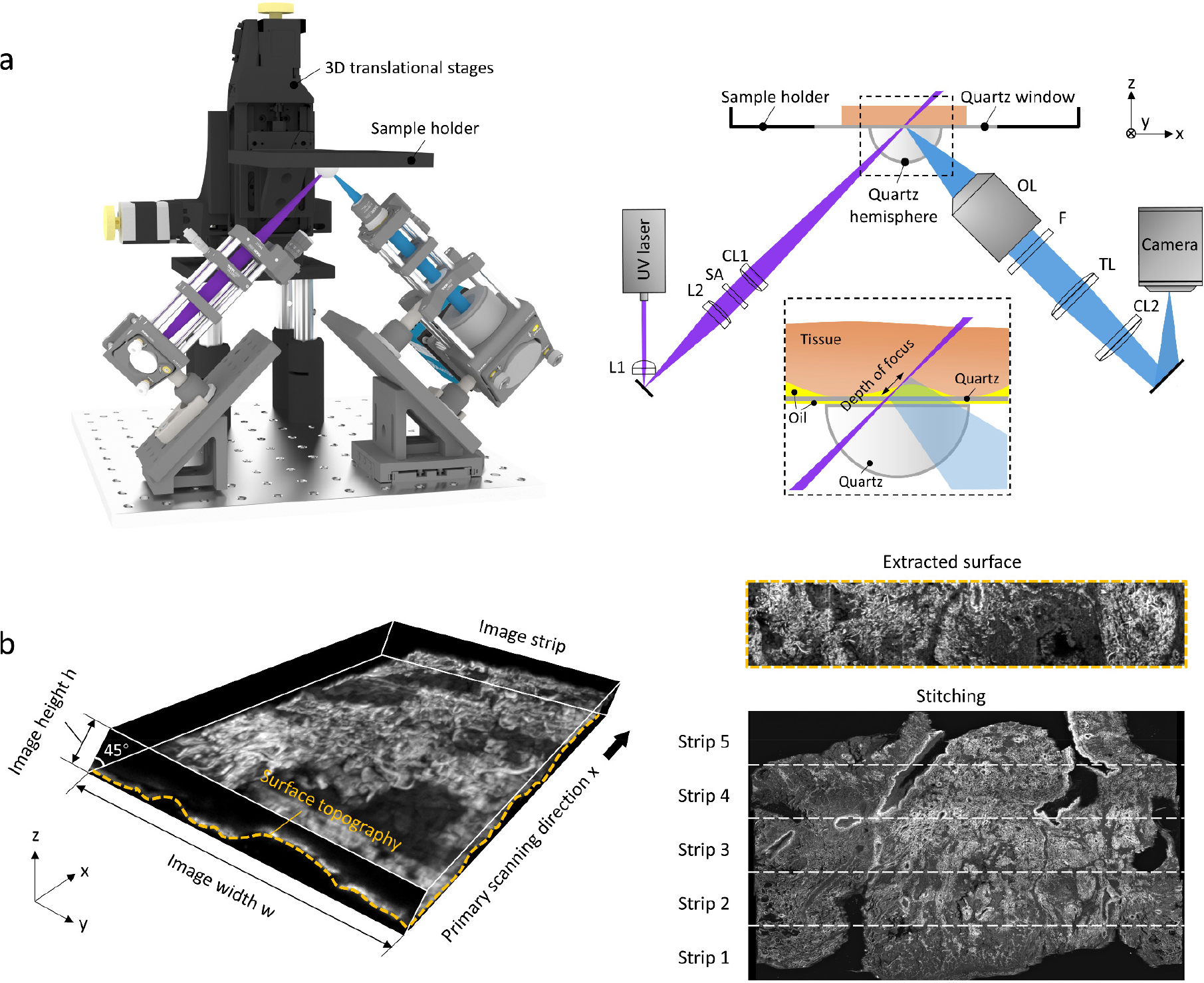
Overview of MUSI imaging. **a**, Schematic of the MUSI system. **b**, Image processing pipeline for MUSI imaging. F: filter; L: lens; SA: slit aperture; CL: cylindrical lens; OL: objective lens; TL: tube lens.

### 2.4. Image processing and data analysis

Figure 1b demonstrates the image processing pipeline for MUSI. The recorded images within an image strip are loaded as a stack (data volume) into MATLAB (MathWorks, Inc) for processing. Note that the tissue geometry reconstructed by the raw data volume will be distorted since the light sheet images are recorded at 45° with respect to the tissue surface. To correct this distortion, the raw data volume is sheared by 45° in the x-z plane to create a trapezoidal data volume [9], [10]. After that, an extended DOF algorithm is implemented on the sheared data volume through a Fiji plugin [29] to extract the intact tissue surface of this image strip. This procedure is repeated until all the image stripes are processed. Then the extracted surfaces of adjacent image stripes are registered and stitched by the grid-stitching plugin in Fiji [30] to produce a large-area tissue image. The processing is jointly completed in MATLAB and Fiji through micro scripts. The algorithm is run on a workstation with a Core i9-10980XE CPU @ 8×32GB RAM, and 4 NVIDIA GEFORCE RTX 3090 GPUs, which takes ∼15 min to process 1-cm^2^ tissue area with a data size of ∼40 GB. To extract the distributions of nuclear features, the MUSI images are segmented and binarized to acquire the cross-sectional area and centroid of each cell nucleus. With the localized center positions of cell nuclei, the intercellular distance is calculated to be the shortest adjacent distance to a neighboring cell nucleus.

## 3. Results

The spatial resolution of MUSI is measured by imaging agarose-embedded sub-diffraction fluorescence beads (B200, 200-nm-diameter, *λ*_em_ = 445 nm, Thermo Fisher Scientific Inc.) across a volume of 0.5 × 1 × 0.05 mm^3^ (Fig. S2a). The full-width at half-maximum (FWHM) of the measured point-spread functions (PSFs) are varied with spatial positions. It is shown that the PSF will deteriorate along the propagation of the light sheet (Fig. S2c,d), and this is more evident at the edge of the imaging FOV (Fig. S2e). The FWHM of the PSF located at the beam waist and center of the imaging FOV (PSF_i_, Fig. S2c) is measured to be ∼2 μm (lateral) and ∼3 μm (axial), which is sufficient to reveal subcellular features (e.g., cell nuclei) and produce an optical section instead of a physical section in slide-based FFPE histology.

### 3.1. *Ex-vivo* imaging of normal mouse tissues

The formalin-fixed mouse brain and kidney tissues are imaged to validate the performance of MUSI initially (Fig. S3), since the fixed tissues are less deformed than fresh and soft tissues during imaging. The tissues are fixed at room temperature for 24 hours and manually sectioned into 3-to 5-mm-thick tissue slabs for imaging, and subsequently histologically processed to obtain the corresponding H&E-stained images for comparison. The cell nuclei located at the hippocampus (Fig. S3b), olfactory area (Fig. S3c), and isocortex (Fig. S3d) in the mouse brain are individually visualized with a negative contrast in the MUSI images, showing high accordance with the corresponding H&E-stained images. Besides, MUSI provides well-characterized structures in the mouse kidney such as glomerular capsules (Fig. S3f) and renal tubules (Fig. S3g), in which densely-packed cell nuclei can be clearly identified. The functional layers of the cortex, outer medulla, and inner medulla in the kidney are well recognized by MUSI based on intensity variations.

Figure 2 further demonstrates the potential of MUSI for label-free and non-destructive imaging of fresh and hydrated tissues. The freshly excised mouse tissues, including the liver (Fig. 2a–c), brain (Fig. 2e–g), and kidney (Fig. 2h–j), are manually sectioned into thick tissue slabs for imaging, after which the tissues are histologically processed to generate the corresponding H&E-stained images for comparison. Note that the FFPE thin slice is not able to exactly replicate the surface imaged by MUSI due to the difference in imaging thickness and tissue processing. During experiments, an image height of 100 μm is sufficient to accommodate surface irregularities presented on the specimens since fresh tissues can sit relatively flattened against the sample holder. We observed that the UV penetration depth typically varies between 5∼30 μm, depending on different tissue architectures and degrees of tissue scattering. For instance, UV penetrates differently in regions with densely-packed cell nuclei (e.g., primary tumors), lipid droplets (e.g., subcutaneous tissues), or large intercellular spaces (e.g., lung alveoli). The morphology of hepatocytes in mouse liver is characterized at a depth of 10-μm below the tissue surface, showing great accordance with the H&E-stained images despite that the sinusoidal capillaries are less identified by MUSI (Fig. 2b). The nuclear features, such as cross-sectional areas and intercellular distances, are extracted from both MUSI and FFPE histology for comparison. The statistical results (Fig. 2d), which are calculated from 50 hepatocytes selected from both MUSI and H&E-stained images in Fig. 2c, suggest that the cellular features extracted by MUSI and the clinical standard method agree fairly well. In the mouse cerebellum, a clear separation between the molecular layer and granular layer with the intermediate Purkinje cells can be visualized in both MUSI and H&E-stained images (Fig. 2f,g). Similarly, the features of renal tubules (Fig. 2i) and glomerulus (Fig. 2j) in the kidney revealed by MUSI are remarkably similar to that by conventional FFPE histology, despite that the Bowman’s space in renal corpuscles is less visible in the MUSI image.

**Figure 2.**
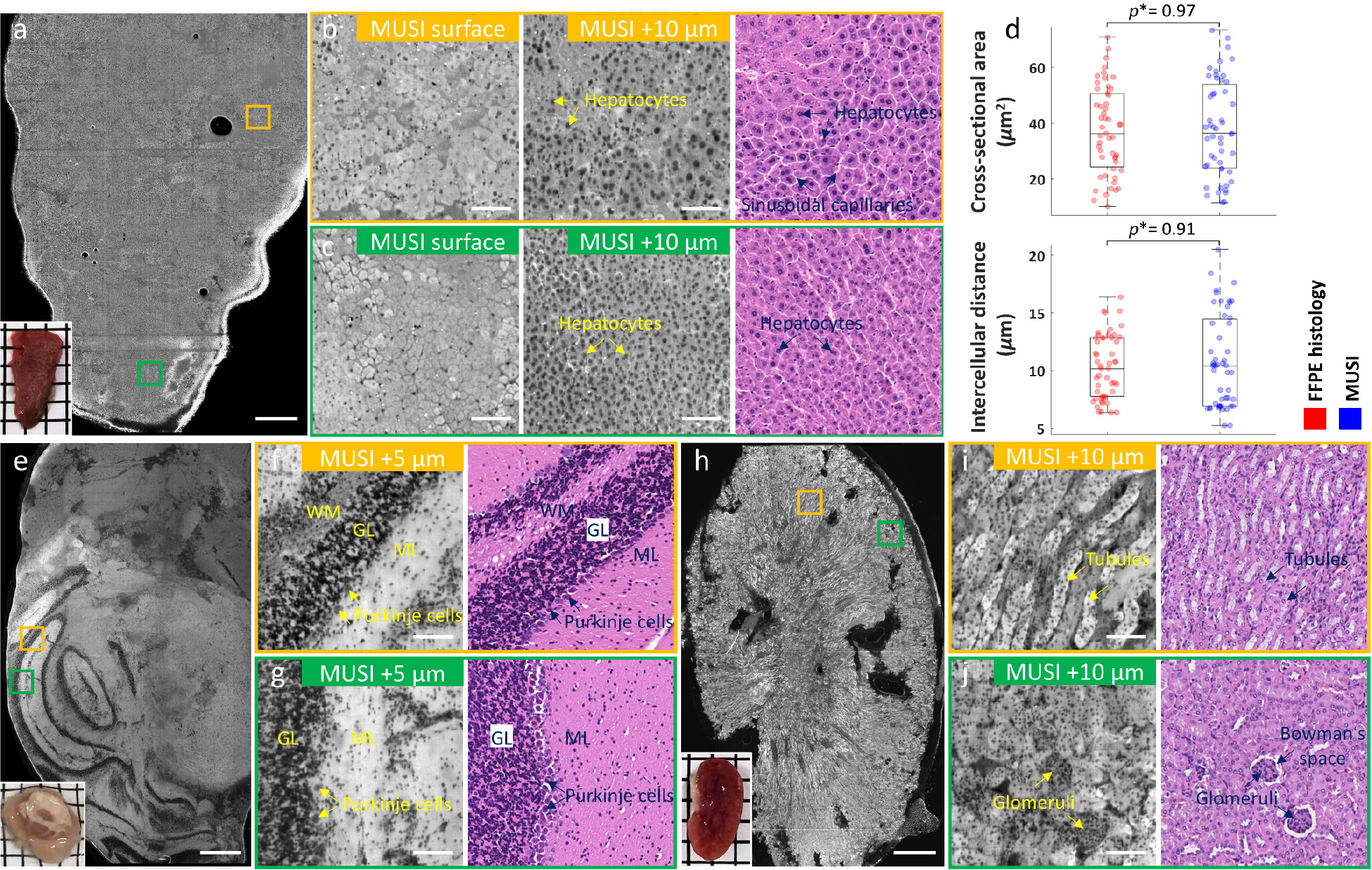
*Ex-vivo* imaging of normal mouse tissues. **a**, MUSI image of a fresh mouse liver, inset at the bottom left shows the photograph of the specimen. **b**,**c**, Zoomed-in MUSI and the corresponding H&E-stained images of the orange and green solid regions marked in a, respectively. **d**, Distributions of nuclear features extracted from c. Wilcoxon rank-sum testing is carried out across groups with *n* = 50 for each distribution. The significance is defined as *p\** ≤ 0.05 in all cases. **e**, MUSI image of a fresh mouse brain, inset at the bottom left shows the photograph of the specimen. **f**,**g**, Zoomed-in MUSI and the corresponding H&E-stained images of the orange and green solid regions marked in e, respectively. **h**, MUSI image of a fresh mouse kidney, inset at the bottom left shows the photograph of the specimen. **i**,**j**, Zoomed-in MUSI and the corresponding H&E-stained images of the orange and green solid regions marked in h, respectively. Scale bars: 1 mm (a,e,h), 100 μm (the remaining). GL: granule layer; ML: molecular layer; WM: white matter.

### 3.2. *Ex-vivo* imaging of cancerous mouse tissues

To further explore the potential of MUSI for rapid screening of cancerous tissues, freshly excised mouse tissues with metastatic human breast cancer, including mouse skin (Fig. 3a–c), lung (Fig. 3d–g), and spleen (Fig. 3h–j), are imaged by MUSI for validation. The MUSI image of a primary tumor (Fig. 3a–c), which is developed through subcutaneous injection and obtained in the skin, is in good accordance with the H&E-stained images in which the densely-packed cancer cells are spreading across the whole slide. The sarcomatoid feature shown in Fig. 3b indicates the invasiveness of the cancer cell line. In addition, it is observed that the mouse lung is invaded by a large number of poorly-differentiated breast tumor cells (Fig. 3e), and the alveolar spaces are spatially compressed by the infiltrating lymphocytes, which will obstruct the normal gaseous exchange in the lung (Fig. 3f). Figure 3g demonstrates a mixture of metastatic tumor cells trying to invade and spread to other parts of the body through the periphery of blood vessels, and immune cells which try to stop the invasion are also observed. Besides, extensive fibrosis can be found in the mouse spleen with splenomegaly (i.e., enlargement of the spleen) (Fig. 3h), which may be related to neoplasm or increased immunologic activity. Disruption of the functional parenchyma, namely red pulp and white pulp, is shown in Fig. 3i, where the morphologically distinctive compartments are indistinguishable and without the presence of follicles and germinal centers. The necrotic cells and connective tissues shown in Fig. 3j further validate the disruptive power of the tumor toward the mouse organs.

**Figure 3.**
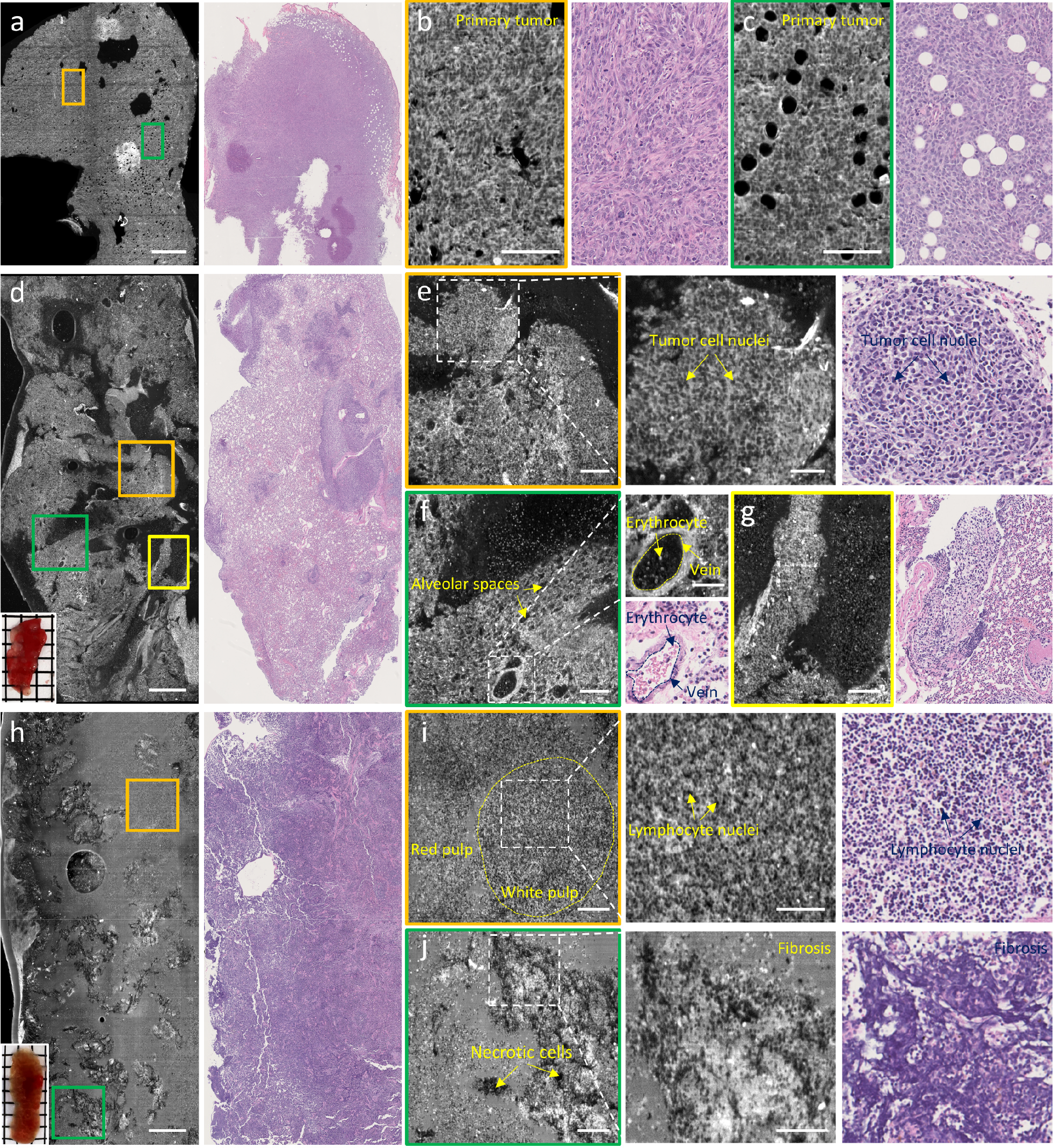
*Ex-vivo* imaging of cancerous mouse tissues. **a**, MUSI and clinical standard image of a primary tumor obtained in mouse skin, inset at the bottom left shows the photograph of the specimen. **b**,**c**, Zoomed-in MUSI and the corresponding H&E-stained images of the orange and green solid regions marked in a, respectively. **d**, MUSI and clinical standard image of a fresh mouse lung with cancer metastasis, inset at the bottom left shows the photograph of the specimen. **e**–**g**, Zoomed-in MUSI and the corresponding H&E-stained images of the orange, green, and yellow solid regions marked in d, respectively. **h**, MUSI and clinical standard image of a fresh mouse spleen with splenomegaly, inset at the bottom left shows the photograph of the specimen. **i**,**j**, Zoomed-in MUSI and the corresponding H&E-stained images of the orange and green solid regions marked in h, respectively. Scale bars: 1 mm (a,d,h), 200 μm (b,c,e–g,i,j), 100 μm (the remaining).

### 3.3. *Ex-vivo* imaging of human lung adenocarcinoma tissues

Adenocarcinoma is the most common type of NSCLC, which typically evolves from mucosal glands and occurs in the lung periphery. Lung adenocarcinomas are usually developed with a mixture of histologic subtypes, including lepidic, acinar, papillary, micropapillary, and solid. The lung cancer specimens (*n* = 15) were obtained from patients who were treated with lung lobectomy with informed consent. The tissues are manually sectioned and imaged by MUSI, and subsequently histologically processed to generate the clinical standard images for comparison. Multiple H&E-stained slices are prepared for each specimen tissue, and the slice that is the most representative of the MUSI image is presented. A partial collection of the imaged specimen is shown in Fig. S4. Figure 4 demonstrates several representative cases of different adenocarcinoma subtypes. Figure 4a showcases a specimen with acinar-predominant adenocarcinoma, where the irregular-shaped glands in a fibrotic stroma can be identified by MUSI (Fig. 4b,c). It is observed that some glands are arranged as solid clusters of tumor cells with a less recognizable lumen (Fig. 4c). Tissue fragments generated through tumor cell breakup can be easily distinguished by MUSI with a much higher fluorescence intensity (Fig. 4d). In comparison, the growth patterns of papillary-or micropapillary-predominant adenocarcinomas are different from that in acinar-type. Figure 4e shows a papillary-predominant lung adenocarcinoma specimen with a positive margin which outlines a clear interface between the normal and cancer regions (denoted by the dotted lines in Fig. 4e). The nuclei of tumor cells are clearly identified at a depth of 10-μm below the tissue surface (Fig. 4f). The finger-like papillary architecture with tumor cells lining the surface of branching fibrovascular cores is revealed in both MUSI and H&E-stained images (Fig. 4g). In comparison, pulmonary alveolus with large air spaces are well characterized in the normal lung tissue (Fig. 4h). Figure 4i demonstrates a pathologically confirmed case of micropapillary-predominant adenocarcinoma. The micropapillary clusters are found floating and dissociative within alveolar spaces with a lack of fibrovascular cores (Fig. 4j,k). Besides, Fig. 4l showcases a region with a large number of tumor-infiltrating lymphocytes and residual anthracosis pigments.

**Figure 4.**
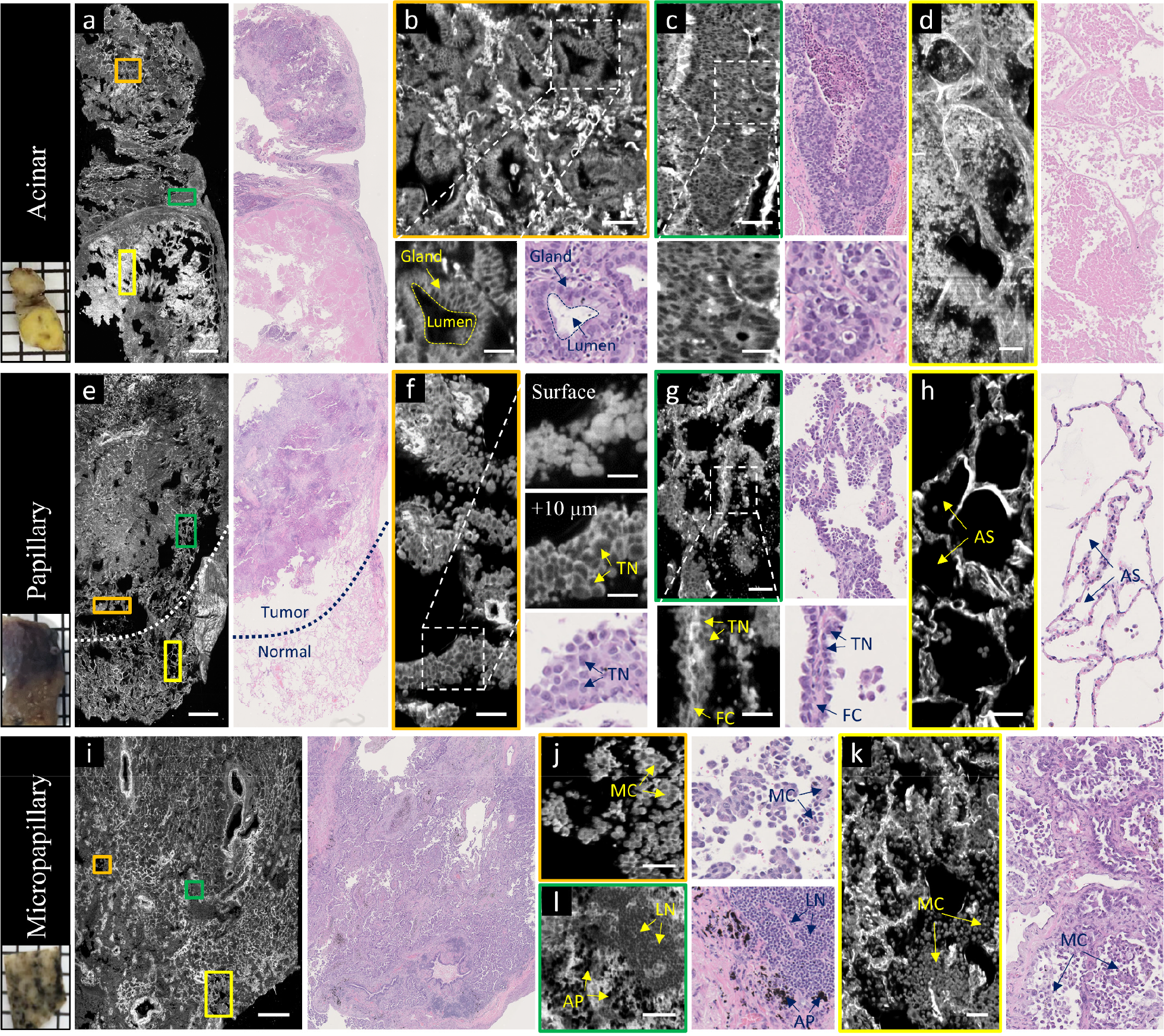
*Ex-vivo* imaging of human lung adenocarcinoma specimens. **a**, MUSI and clinical standard image of a lung specimen with acinar-predominant adenocarcinoma, inset at the bottom left shows the photograph of the specimen. **b**–**d**, Zoomed-in MUSI and the corresponding H&E-stained images of the orange, green, and yellow solid regions marked in a, respectively. **e**, MUSI and clinical standard image of a lung specimen with papillary-predominant adenocarcinoma, inset at the bottom left shows the photograph of the specimen. **f**–**h**, Zoomed-in MUSI and the corresponding H&E-stained images of the orange, green, and yellow solid regions marked in e, respectively. **i**, MUSI and clinical standard image of a lung specimen with micropapillary-predominant adenocarcinoma, inset at the bottom left shows the photograph of the specimen. **j**–**l**, Zoomed-in MUSI and the corresponding H&E-stained images of the orange, yellow, and green solid regions marked in i, respectively. Scale bars: 1 mm (a,e,i), 100 μm (b–d,f–h,j–l), 50 μm (the remaining). TN: tumor cell nuclei; LN: lymphocyte nuclei; FC: fibrovascular core; AS: alveolar space; MC: micropapillary cluster; AP: anthracosis pigment.

### 3.4. *Ex-vivo* imaging of other human tissues

The wide applicability of MUSI is also validated by other types of tissues such as human skin (Fig. S5a–d) and human brain (Fig. S5e–g). These tissues are considered as leftover tissue, i.e., it represents a portion of a collected specimen that is not needed for the diagnosis and treatment of the patient. It is shown that a variety of histologic features can be simultaneously visualized by MUSI with deep-UV excitation. For instance, anatomic structures in human skin, including the sudoriferous gland (Fig. S5b), erythrocyte-filled arterial lumen (Fig. S5c), and adipose tissue (Fig. S5d), are well characterized by MUSI in a label-free and non-destructive manner. These features share great similarities with that in slide-based FFPE histology. In addition, cerebellar granule cells are resolved individually in the human brain (Fig. S5f), and neuroglial cells can be visualized within a depth range of 30 μm (Fig. S5g).

### 3.5. *In-vivo* imaging of intact mouse tissues

In addition to *ex-vivo* imaging of resected tissues, we further explore the potential of MUSI for rapid *in-vivo* imaging of intact tissues. Firstly we imaged several freshly excised organs without cutting open the specimens to verify that histologic features can be extracted from the surface of intact tissues (Fig. S6). Slight pressure is applied from the top of these samples to make sure they are sufficiently contacted with the sample holder during imaging. After imaging, multiple H&E-stained thin slices are prepared for each specimen, and the slice that is the most representative of the imaged surface by MUSI is shown for comparison. We found that a variety of anatomic structures, including vessels and cell nuclei in the brain cortex (Fig. S6a), renal tubules in the kidney (Fig. S6b), muscle fibers in the anterolateral thigh (Fig. S6c), and filiform papillae in dorsum tongue (Fig. S6d), are clearly revealed and in good accordance with the conventional slide-based histology.

Figure 5 demonstrates the MUSI’s capacity in label-free *in-vivo* imaging of mouse brain and kidney tissues. With a rapid acquisition speed of 250 frames/s, anatomic features can be captured without obvious breath-induced motion artifacts. The energy fluence measured at the imaging plane is ∼2 mJ/cm^2^, which is below the safety threshold regulated by the American Conference of Governmental Industrial Hygienists (ACGIH^®^) [31]. Figure 5c shows an area in the brain that contains histologic features from both the cerebral cortex and pia mater. The cell nuclei located at the surface of the brain cortex can be resolved individually with a negative contrast (Fig. 5d). The pia mater, which is the innermost layer of the meninges that clings tightly to the brain, can be also identified (Fig. 5e). We can observe from both MUSI and H&E-stained images that the pia mater is rich vascularized and consists of a thin layer of squamous epithelial cell. Figure 5f shows a surrounding area in the dorsum ear that is filled with epidermal keratinocytes (Fig. 5g). While in the mouse kidney, due to the highly fluorescent cytoplasmic lipofuscin and urinary cast material, the renal tubules at the outermost cortex can be visualized with a high signal-to-noise ratio (SNR) (Fig. 5j,k). In addition, other morphological structures such as adipose tissue in perinephric fat (Fig. 5l) and connective tissues in the lumbodorsal fascia (Fig. 5m) can be also identified. Although MUSI cannot provide 1:1 imaging to conventional histology correlates due to the difference in imaging depth and tissue processing, the tissue morphology remains remarkably similar from the gross to the subcellular level. The ability of MUSI to image the intact tissues non-destructively can potentially find applications that preclude invasive biopsy procedures. Compared with recently proposed imaging platforms for label-free *in-vivo* tissue pathology, such as SLAM [32], MediSCAPE [33], and LC-OCT [34], MUSI may have limited access to deep tissue information. However, MUSI significantly strengthens the contrast between nucleic acid and extracellular matrix with deep-UV absorption, thus showing higher consistency with the histochemical staining in routine pathological practice. Due to this reseason, our images can be easily virtually stained to mimic the appearance of the H&E-stained images via deep learning-based style transfer frameworks [25], [35] (Fig. S7). This will ensure an easy adaption by pathologists and produce a great clinical impact.

**Figure 5.**
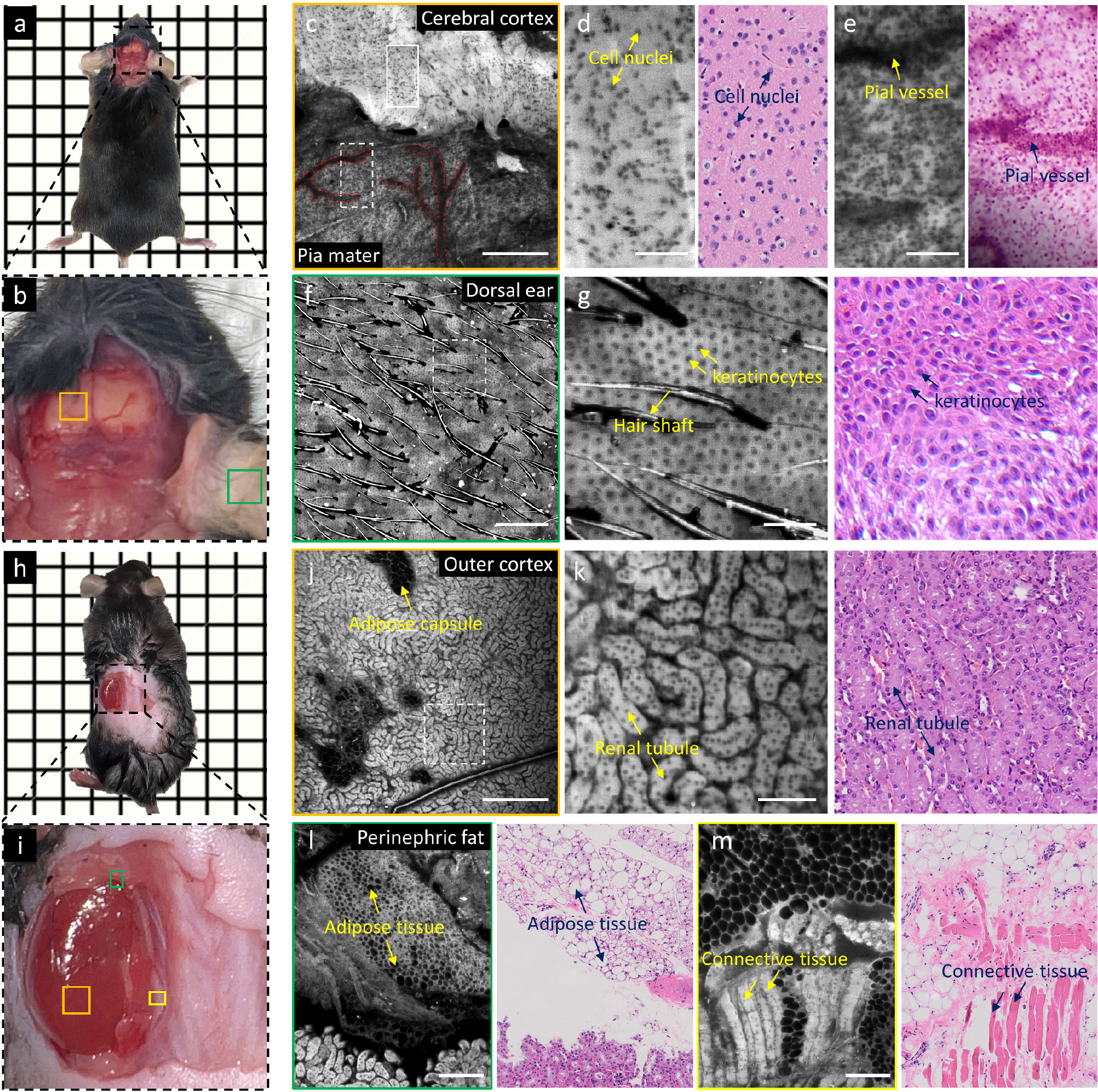
*In-vivo* imaging of intact mouse tissues. **a**,**b**, Photograph of the imaged brain with parietal craniotomy. **c**, Zoomed-in MUSI image of the orange solid region marked in b. Pial vessels are depicted by red dotted lines. **d**,**e**, Zoomed-in MUSI and the corresponding H&E-stained images of the white solid and dashed regions marked in c, respectively. **f**, Zoomed-in MUSI image of the green solid region marked in b. **g**, Zoomed-in MUSI and the corresponding H&E-stained image of the white dashed region marked in f. **h**,**i**, Photograph of the imaged kidney with lumbar nephrectomy. **j**, Zoomed-in MUSI image of the orange solid region marked in i. **k**, Zoomed-in MUSI and the corresponding H&E-stained image of the white dashed regions marked in j. **l**,**m**, Zoomed-in MUSI and the corresponding H&E-stained images of the green and yellow solid regions marked in i, respectively. Scale bars: 500 μm (c,f,j), 200 μm (l,m), 100 μm (the remaining).

## Conclusion

MUSI enables rapid *ex*-or *in*-*vivo* assessment of fresh and unprocessed tissues in a label-free and non-destructive manner, holding great promise to revolutionize the current clinical practice in tissue pathology. However, there are still some deviations between the MUSI and conventional FFPE histology. First, anatomic structures, such as sinusoidal capillaries in mouse liver and Bowman’s space in mouse kidney, are well identified by FFPE histology but hardly visualized by MUSI (Fig. 2b,j). This is probably due to the grossly-cut soft tissues would be deformed when being flattened and placed on the sample holder. Second, fibrous structures, such as fibrotic stroma in human lungs (Fig. 4b,k) and elastic membrane in the artery (Fig. S5c), are better visualized in MUSI than that in H&E-stained images. This is because these structural proteins present a high quantum yield with deep-UV excitation while eosin exhibits a similar affinity across the cytoplasm. Third, the nucleoli features are less recognizable in the MUSI images (e.g., Fig. 5d,g and Fig. S3d), which is likely because the fluorescence property of nucleoli in the detected spectral range is not chemically identical to the histological stains.

Intrinsic fluorescence with deep-UV excitation naturally forms a contrast mechanism for label-free tissue imaging. However, the fluorescence properties of endogenous fluorophores, such as emission maximum and quantum yields, are highly related to tissue phenotypes and disease status, causing obvious intensity variations in the MUSI images. For instance, the spleen, the largest mass of lymphoid organs, usually presents the least fluorescence among other types of tissues. In cancerous human lungs, solid-predominant adenocarcinoma with a lack of fibrotic stroma shows a relatively weak intensity compared with other histologic subtypes. In addition, we observed that the overall intensity will be increased with continuous UV radiation. The combination of MUSI imaging and autofluorescence spectroscopy holds great promise to further increase the accuracy in the diagnosis of lung cancer, as some metabolic enzymes (e.g., NADH and FAD) are highly associated with cellular changes and can effectively serve as a tumor-specific biomarker [26]. A variety of anatomic features, such as cell nuclei and blood vessels, can be simultaneously visualized by MUSI through non-specific excitation, which potentially poses a challenge for the accurate segmentation of the clinically relevant features. We believe that this limitation can be mitigated by using advanced neural networks which would enable a more faithful tissue analysis [36].

MUSI is currently in an early stage of development. The system can be further optimized to achieve higher imaging performance for wide clinical applications. The current system can achieve an imaging speed of 0.5 mm^2^/s, which is predominantly restricted by the weak intrinsic fluorescence of tissues. For each raw image in the sequence of primary scanning, an integration time of 4 ms is required to maintain an acceptable SNR under an illumination power of 2 mW, such that the camera is only allowed to operate at a framerate of 250 fps. The imaging speed is expected to be increased by an order of magnitude with a high-power UV illumination source that could reduce the exposure time to a few hundred microseconds. In this case, the imaging speed will be determined and limited by the degrees of tissue irregularities. However, *in-vivo* applications are prohibited since the strong UV radiation may cause damage to internal organs. With MUSI’s dual-axis configuration, imaging parameters, such as DOF and lateral/axial resolution, can be optimized separately. In the illumination arm, the trade-off between the length of a Gaussian light sheet (i.e., DOF, 2*Z*_R_) and its thickness (i.e., 2*w*_0_) is inevitable due to the diffraction of light. In practice, the DOF could be largely extended without the compromise of axial resolution by using the light sheet generated through rapid scanning of non-diffracting beams [37], [38]. This can further improve the system tolerance for large irregular surfaces. In the detection arm, an in-plane resolution of ∼2 μm is still not sufficient for high-resolution interrogation of certain diagnostic features (e.g., nucleoli). To satisfy the clinical needs for multi-scale imaging, a switchable set of objective lenses with resolutions from sub-micron to macroscopic scales can be implemented in the current system. In addition, the development from a bentch-top to a probe-based handheld MUSI device can greatly increase the flexibility of operation, promoting the applications of intraprocedural biopsy guidance. As an assistive imaging platform, MUSI could potentially cooperate with other surgical navigation techniques such as ultrasound and computed tomography [39] to enable precise localization of cancer margins and residual lesions. This could help minimize surgical complications and ultimately improve the outcome of the surgery. Moreover, augmented reality can be integrated into these image-guided procedures to help surgeons with improved accuracy and safety [40].

In summary, the proposed MUSI method enables label-free and non-destructive imaging of fresh and unprocessed tissues, allowing centimeter-scale surgical specimens to be screened with a subcellular resolution within 3 minutes. It is experimentally demonstrated that MUSI can provide diagnostically important features to differentiate between different subtypes of human lung adenocarcinomas, holding great promise to streamline the current workflow in surgical pathology. As a proof-of-concept study, this work is limited by a small number of specimens (*n* = 15). Large-scale clinical trials should be carried out as follow-up work to quantify the diagnostic metrics (i.e., sensitivity and specificity) and study the inter-patient variations. In addition, computer-aided diagnosis can be also incorporated with MUSI to further enhance the current clinical practice.

## Data Availability

All data involved in this work, including raw/processed images provided in the paper, are available from the corresponding author upon request.

## Code availability

The customized MATLAB code for image processing is available from the corresponding author upon request.

## Acknowledgments

The Translational and Advanced Bioimaging Laboratory (TAB-Lab) at HKUST acknowledges the support of the Hong Kong Innovation and Technology Commission (ITS/036/19); Research Grants Council of the Hong Kong Special Administrative Region (16208620 and 26203619).

## Author contributions

Y. Z. and T. T. W. W. conceived of the study. Y. Z., B. H., and L. K. built the imaging system. Y. Z., B. H., and V. T. C. T. prepared the specimens involved in this study. Y. Z., B. H., and J. W. performed imaging experiments. C. T. K. L. performed histological staining. Y. Z. processed the data. Y. Z. and T. T. W. W. wrote the manuscript. T. T. W. W. supervised the whole study.

